# Variance component estimates, phenotypic characterization, and genetic evaluation of bovine congestive heart failure in commercial feeder cattle

**DOI:** 10.1101/2023.01.19.524739

**Authors:** Justin W. Buchanan, Lex E. Flagel, Michael D. MacNeil, Ashley R. Nilles, Jesse L. Hoff, Joseph K. Pickrell, Randall C. Raymond

## Abstract

The increasing incidence of bovine congestive heart failure (BCHF) in feedlot cattle poses a significant challenge to the beef industry due to economic loss, reduced performance, and reduced animal welfare attributed to cardiac insufficiency in growing animals. Changes to cardiac morphology as well as abnormal pulmonary arterial pressure (PAP) in purebred cattle of mostly Angus ancestry have been extensively characterized over the past 30 years. However, congestive heart failure affecting cattle late in the feeding period has been an increasing problem over time and tools are needed for the industry to address the rate of mortality in the feedlot. At harvest, a population of 32,763 commercial fed cattle was phenotyped for cardiac morphology with associated production data collected from feedlot processing to harvest at a single feedlot and packing plant in the Pacific Northwest. A sub-population of 5,001 individuals were selected for low-pass genotyping to estimate variance components and genetic correlations between heart score and the production traits observed during the feeding period. At harvest, the incidence of a heart score of 4 or 5 in this population was approximately 4.14%, indicating a significant proportion of feeder cattle are at an increased risk of cardiac mortality before harvest. The heart scores were also significantly and positively correlated with the percentage Angus ancestry observed by genomic breed percentage analysis. The heritability of heart score measured as a binary (scores 1 and 2 = 0, scores 4 and 5 = 1) trait was 0.356 in this population, which indicates the development of a selection tool to reduce the risk of congestive heart failure in the form of an EPD (expected progeny differences) is feasible. Genetic correlations of heart score with growth traits and feed intake were moderate and positive (0.289 to 0.460). Genetic correlations between heart score and backfat and marbling score were -0.120 and -0.108, respectively. Significant genetic correlation to traits of high economic importance in existing selection indexes may explain the increased rate of congestive heart failure observed over time. These results indicate there is potential to implement heart score observed at harvest as a phenotype under selection in genetic evaluation in order to reduce feedlot mortality due to cardiac insufficiency and improve overall cardiopulmonary health in feeder cattle.

## 1 Introduction

Bovine congestive heart failure (BCHF) observed in feedlot cattle is responsible for significant and increasing economic loss to the feedlot industry, with the largest losses occurring during the latter part of the finishing period (Neary et al., 2015a; Johnson et al., 2021). This condition has been characterized in a variety of studies and is commonly referred to as congestive heart failure, right-sided heart failure, cor pulmonale, or noninfectious heart disease. These conditions generally describe morbidity due to changes in cardiopulmonary function and morphology that can result in reduced performance and in some cases mortality prior to harvest. Data compiled from 22 commercial feedlots indicates the rate of mortality from non-infectious heart disease is approximately 4% of all feedlot deaths (Johnson et al., 2021). However, the consistency of post-mortem diagnosis of cardiac related deaths among feedlot personnel is unknown. Multiple studies have identified underlying pathologies associated with congestive heart failure cases including pulmonary hypertension, right-ventricular hypertrophy, ventricular fibrosis, diastolic dysfunction, vascular remodeling, and abdominal edema (Buczinski et al., 2010; Krafsur et al., 2018; Moxley et al., 2019). It has been proposed that end stage heart failure in fattening cattle that are otherwise healthy is the result of a cascade of cardiopulmonary changes beginning with pulmonary hypertension, ventricular fibrosis and stiffening of the myocardium, diastolic dysfunction, and ultimately severe ventricular remodeling leading to heart failure and mortality (Krafsur et al., 2018). Changes in cardiopulmonary morphology linked to the progression of BCHF can be visually identified upon inspection of the viscera in cattle that reach harvest, with the most severe cases presenting a flaccid and rounded or blunt shape of the heart (Heffernan et al., 2020). It is hypothesized that this cascade of changes is the direct result of increased rates of growth and adiposity that drive hypoxia and pulmonary hypertension, and that these underlying issues have been inadvertently selected for in modern feedlot cattle (Krafsur et al., 2018; Norouzirad, et al., 2017).

High altitude disease, or brisket disease, is a closely related complex of symptoms that may result in congestive heart failure associated primarily with pulmonary hypertension observed in cattle at elevations greater than 1600 m (Malherbe et al., 2012; Cockrum et al., 2014; Crawford et al., 2016; Neary et al., 2015a; Jennings et al., 2019). Prevalence of this condition, as well as mortality due to congestive heart failure in feedlot cattle at moderate and low altitudes, has been increasing, especially in the cattle feeding regions of the West and high plains (Neary et al., 2015a; Johnson et al., 2021). Extensive research has been carried out to characterize pulmonary hypertension at altitude, especially in the Angus breed (Shirley et al., 2008; Neary et al., 2015b; Newman et al., 2015; Pauling et al., 2018). Evidence suggests pulmonary arterial pressure (PAP) is at least moderately heritable in purebred populations that were phenotyped for PAP at elevation, and it would be reasonable to expect non-zero genetic variation in a phenotype evaluating morphology of the heart in harvested cattle (Weir et al., 1974; Cockrum et al., 2014; Crawford et al., 2016; Kukor et al., 2021). However, measuring PAP on a large volume of animals over multiple generations requires significant cost and a veterinarian trained in jugular catheterization for invasive blood pressure monitoring (Holt and Callan, 2007). This trait has traditionally been characterized only in purebred and seedstock animals due to the cost and time inputs required for its measurement. The direct genetic correlation between PAP and cardiopulmonary health and animal performance in commercial feeder cattle is currently unknown. These factors have limited the usefulness of the PAP phenotype in genetic evaluation systems and resulted in the collection of relatively few phenotypes in comparison to routinely measured traits such as early-in-life weights, and carcass characteristics measured using ultrasound. Collection of phenotype data evaluating cardiopulmonary health in commercial cattle at harvest would allow the collection of a large volume of data and enable subsequent development of better selection tools to improve the ability to manage this trait in the seedstock sector.

The objectives of this study were to: 1) characterize the distribution of heart scores in commercial feeder cattle originating from a single large feedlot in the Pacific Northwest, 2) analyze breed effects, production traits, and carcass characteristics associated with heart scores observed at harvest, 3) estimate variance components and genetic correlations of BCHF and associated production traits, and 4) develop and evaluate a prototype BCHF EPD derived from packing plant phenotypes collected from commercial cattle.

## 2 Methods

### 2.1 Cattle Enrollment and Data Collection

All cattle enrolled in this analysis were processed at a commercial feedlot in Grand View, Idaho at 756 m of elevation (N = 32,763 head). Cattle were sourced from a variety of operations from the West and Pacific Northwest and include commercial cow-calf and ranch origins as well as beef-on-dairy and dairy breed origins. Upon feedlot entry, all calves were processed according to standard commercial feedlot procedures. A subset of calves were assigned a unique electronic identification (EID) tag during feedlot processing or tracked using an existing EID tag assigned prior to feedlot arrival. A processing weight, coat-color phenotype or observed breed type, and a DNA sample from tail hair linked to a unique barcode were also captured at processing. All data was recorded using feedlot data collection software (CattleXpert, CattleXpert LLC, Elkhorn, NE; CattleInfo, Simplot Livestock Co., Grand View, ID). Calves were placed in pre-assigned pens at an average stocking rate of 12.8 m^2^/head and offered ad-libitum access to feed and water consistent with commercial feedlot practices. Potato by-products (processing or fry waste) and corn in various forms (flaked, high moisture, or dried distillers’ grains) were the primary source of dietary energy. All animals were weighed when they were re-implanted, and live weight was recorded at harvest to assess average daily gain (ADG kg/day) in the early (processing to between 80 and 120 days on feed), late (120 to 80 days prior to harvest), and full (processing to harvest) portions of the feeding period. All treatment events for morbidity were recorded as well.

All cattle were harvested at CS Beef Packers in Kuna, Idaho. At harvest, a full body weight was captured as well as the EID assigned or recorded at processing. Carcass data was captured including hot carcass weight (HCW), ribeye area (REA), backfat thickness (FAT), marbling score (MARB), and yield grade (YG) utilizing camera image analysis (VBG 2000, e+v Technology GmbH & Co. KG, Oranienburg, Germany). Additionally, trained personnel were positioned at the viscera removal location to capture individual cardiac remodeling scores using a 1-5 heart scoring guide (Figure 1) developed by Heffernan et al. (2020). A heart score of 1 represents a normal heart with a conical shape, well defined apex, and firm muscular structure of the left and right ventricles. As heart score progresses to a 5, the musculature of the heart becomes flaccid, a blunting of the apex is observed, and the left and right ventricles become enlarged and flaccid giving an overall deflated appearance to the structure. For binary statistical analysis individuals with a heart score of 1 or 2 were defined as controls and individuals with a heart score of 4 or 5 were defined as BCHF cases.

**Figure 1.**
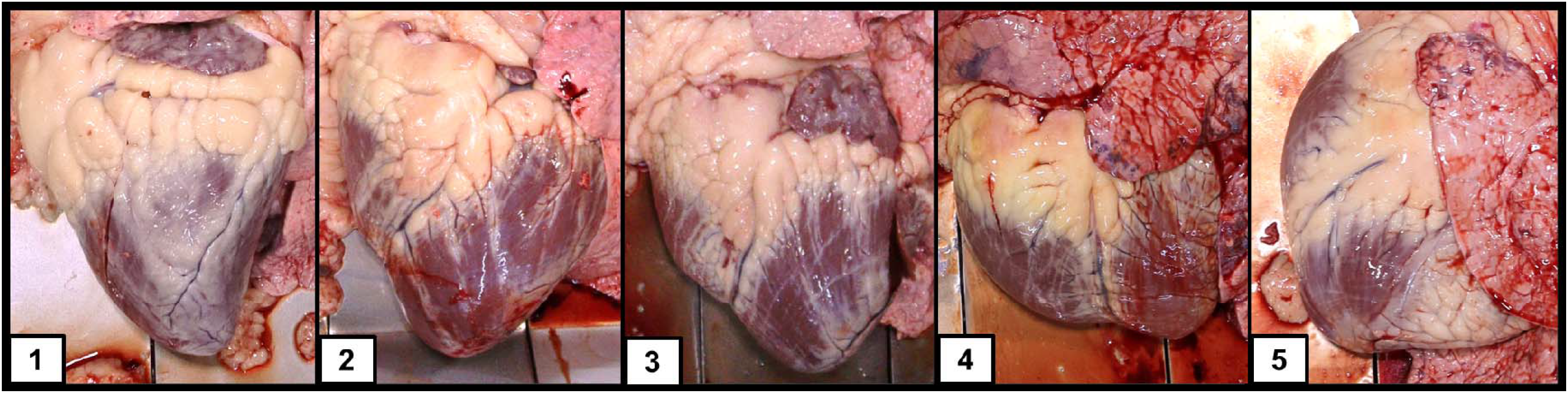
Heart images taken at harvest from growing feeder cattle representative of the 5 categories of heart scores.

A subset of individuals included in this analysis were selected using an emergency harvest identification and shipping procedure developed by Simplot Livestock Co. and implemented by trained feedlot personnel. Horseback pen riders were trained under the supervision of a D.V.M. to identify individual cattle at risk of feedlot mortality by observing intolerance to exercise, elevated respiration rate, increased salivation, abnormal posture, general malaise and/or failure to thrive. Emergency harvest shipments were carried out weekly from approximately May through September and represent a population of cattle at higher risk of exhibiting a heart score greater than one combined with a lighter finished weight at harvest due to early shipment. It is important to note that cattle identified for emergency harvest were not exclusively suspected of congestive heart failure but could be pulled for a variety of other issues including but not limited to chronic lung lesions, kidney failure, pericarditis, or physical injury.

### 2.2 Descriptive statistics

Least square means for phenotypes observed from processing through harvest were generated from regression analysis performed in JMP 12.2.0 (SAS Institute Inc., Cary, NC). Each phenotype was fit as a dependent variable in a regression model with fixed effects of lot (processing groups describing origin), harvest date, and heart score (1 through 5). Least square means were extracted from the model for animals harvested within their original lot assigned at processing. Tukey’s Honest Significant Difference test of multiple comparisons was used to determine differences among least square means (P < 0.05). Animals harvested under emergency shipment protocol were not included in the estimates of least square means since they were harvested earlier than their contemporaries and did not represent groups of harvested animals of the same age or similar origins. Descriptive statistics including the mean and standard deviation for carcass data were calculated separately for animals harvested under emergency shipment protocols since a wide range of days on feed was observed in this population.

### 2.3 Genomic Analysis and Genotype Imputation

A subset of 5,001 individuals were selected for Low-Pass whole genome sequencing (Gencove Inc., New York, NY). These individuals represent two sampling years (2020 and 2021) with 4,639 sampled from year 1 (373 cases and 4,266 controls), and 362 from year 2 (182 cases and 180 controls). Harvest dates and heart score phenotyping occurred from approximately March through October of each sampling year. DNA samples for genotyping were collected either upon feedlot processing from tail hair or at harvest using blood cards taken at the time of cardiac scoring. All individuals with a DNA sample on record and a heart score of 4 or 5 (cases) were selected for Low-Pass genotyping, and a random sample of individuals from the same harvest cohort with a heart score of 1 or 2 were selected as controls.

Low-Pass whole genome sequencing was performed on all 5,001 individuals, resulting in an average sequenced genome coverage of approximately 0.5x per individual (i.e. ∼1.5e9 base pairs). Whole genome imputation of this data was performed using the method described in Snelling et al. (2020), resulting in genotype calls for every individual at 59,204,179 genomic sites. The ∼59M Low-Pass imputed loci were filtered to 497,722 sites by first removing all sites with minor allele frequency < 10% (resulting in approximately 9M sites), and then dividing the genome into 497,722 equally spaced bins and randomly selecting 1 locus per bin to be used for further genomic analyses. Principal component analysis (PCA) was carried out on filtered genotypes using PLINK 2.0 (Chang et al., 2015) with the cow option.

In addition to SNP marker calls generated from Low-Pass genotyping, genomic breed percentages were generated for each individual. To estimate genomic breed percentages, whole genome data was compiled from a set of 734 animals across 13 cattle breeds. A supervised version of the ADMIXTURE model (Alexander et al., 2009) was implemented to assign breed percentages for each individual from these 13 breeds. Generalized breed descriptions were captured at processing based on visual appearance of phenotype along with known breed or breeds of origin, and this generalized category was cross checked with genomic breed percentage. A binary logistic regression of heart score on breed percentage was also carried out to assess the relative risk of a heart score of 4 or 5 associated with the major breed types observed in the population of genotyped individuals.

Variance components were estimated for BCHF and the additional production traits using a binary 2-trait threshold-linear genomic model with a Bayesian approach implemented in THRGIBBS1F90 (Tsuruta and Misztal, 2006). Low-pass genotype calls were filtered down to a set of 36,414 SNP markers that were overlapping with markers on the commercially available GGP Bovine 100K array (Neogen Corporation, Lansing, MI). The underlying distribution for the bivariate linear-threshold animal model is assumed:

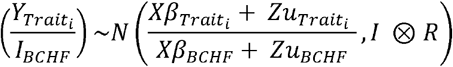

where β are fixed effects including ranch of origin, weaning date, and sex; u are breeding values; X and Z are incidence matrices that link data with respective effects; and R is the residual covariance matrix. The response for BRD was modeled with the following distribution:

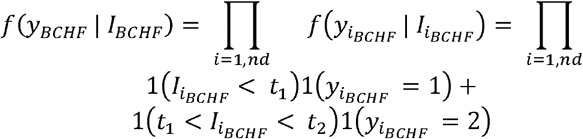

where t1 and t2 are thresholds that define the 2 categories of response. All prior distributions were assumed flat. Within THRGIBBS1F90, a single chain of 50,000 iterations was used to calculate posterior means with the first 10,000 iterations discarded as burn-in and every sample stored for post-analysis. The remaining samples were used to calculate posterior means to estimate variance components and heritability with the program POSTGIBBSF90 (Tsuruta and Misztal, 2006).

### 2.4 Genomic Prediction Model

For EPD estimation the down-sampled sites (497,722) were converted to -1, 0, and 1, corresponding to reference genome homozygote, heterozygote, and alternative allele homozygote, respectively. Phenotype data was converted to cases (heart score ≥ 4) and controls (heart score ≤ 2), and these case/control phenotypes along with the converted genotypes were fitted with a GBLUP model using the rrBLUP package in R (Endelman 2011). To estimate GBLUP model performance on testing/training splits, the Pearson’s correlation between predicted and true case/control BCHF phenotypes was used. For each test split the BCHF phenotypes for that set of individuals were removed from the model, while the remaining training split contained all phenotypes for individuals in the training population.

## 3 Results and Discussion

The cattle phenotyped in this analysis were representative of typical cattle on feed in the Pacific Northwest with ages ranging from approximately 12 to 18 months. Least square means for production and carcass traits for individuals with a heart score phenotype are displayed in Table 1. There were 32,763 individuals harvested without the use of the emergency harvest procedures that had a heart score and carcass data recorded in this dataset. Individuals with a heart score of 4 or 5, which represents end-stage congestive heart failure, were observed at a rate of 4.14% in the overall population. Approximately 39.54% of the population was observed to have some level of cardiac remodeling (heart score 2, 3, 4 or 5). Heart score phenotypes observed at harvest from large populations of feedlot cattle have not been previously reported in the literature. A study by Kukor et al. (2021) reported heart scores from 632 steers and heifers originating from a single feedlot, with 34% of the individuals observed to have a heart score of 3 or greater. However, individuals having a heart score of 5 were not observed in the Kukor et al. (2021) dataset which may be due to the relatively small number of cattle observed or the seasonal severity of BCHF (Neary et al., 2016; Johnson et al., 2021) since heart scores were only recorded for cattle harvested from December to February. Epidemiological studies examining reported feedlot mortality attributable to BCHF indicate the incidence of this trait is increasing over time (Johnson et al., 2021; Neary et al., 2016), however studies observing progression of cardiac morphology in normal harvest lots of feedlot cattle have not been previously reported.

**Table 1.**
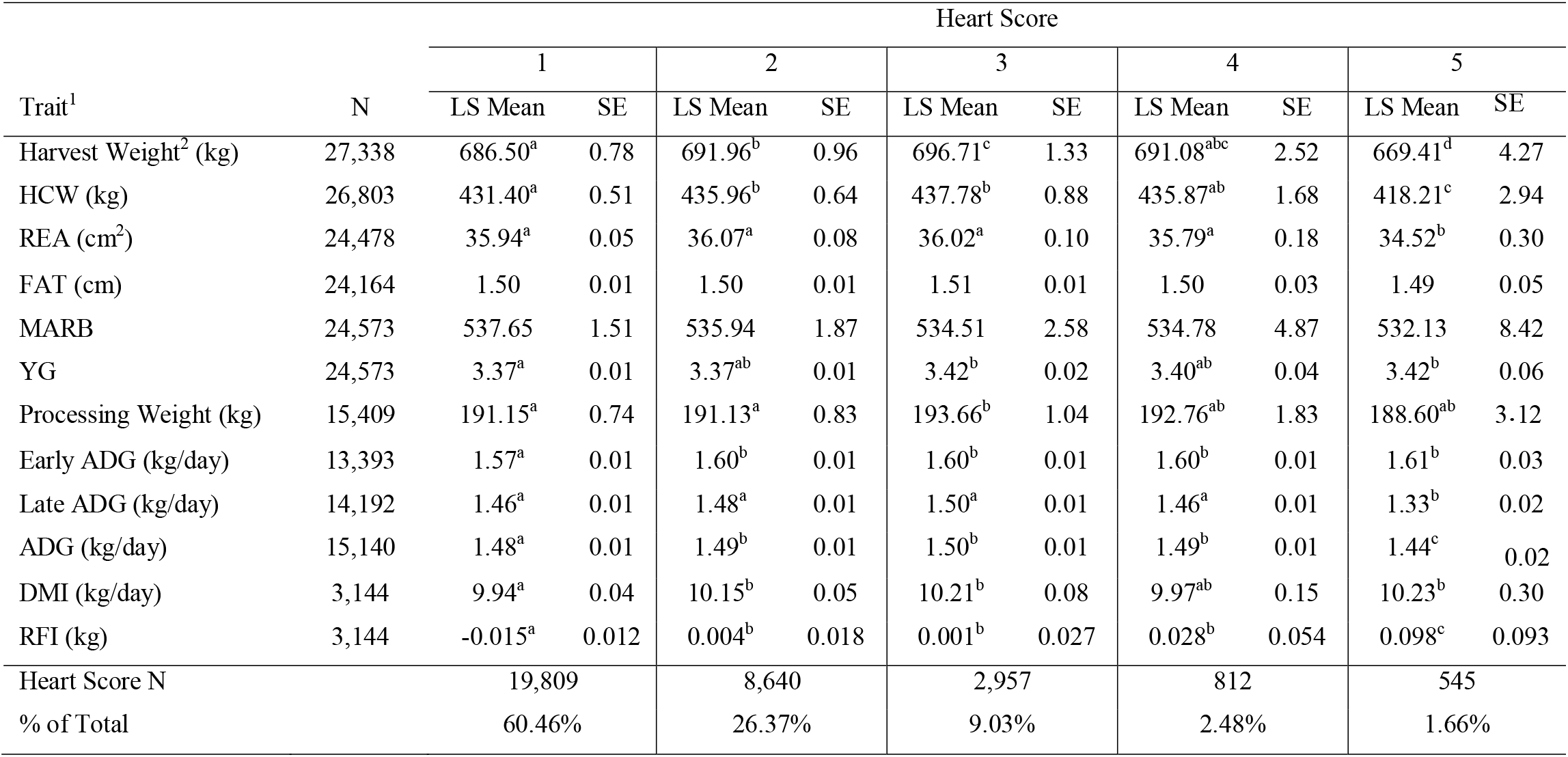
Least square means, standard error, and observed incidence for production and carcass traits for individuals with a heart score of 1 to 5.

Body weight at harvest varied among individuals with different heart scores (Table 1), with the heaviest individuals having a heart score of 2, 3 or 4 (P < 0.05). Individuals with a heart score of 5 at harvest were significantly lighter than animals with any other heart score (P < 0.05). This difference was also observed for HCW (P < 0.05). Processing weight at feedlot entry followed a similar trend as harvest weight, with the heaviest animals displaying a heart score of 3 compared to animals with a heart score of 1 or 2 (P < 0.05). Yield grade, as determined by camera image analysis, differed among heart score categories, with the highest yield grade occurring in animals with a heart score of 3 or 5 compared to animals with a heart score of 1 or 2 (P < 0.05). The lack of evidence of an association between backfat or marbling and heart score is somewhat unexpected given the logical association between fatness, hypoxia, pulmonary hypertension and the proposed mechanism of congestive heart failure in humans (Ebong et al., 2014). However, investigation of heart failure outcomes in humans has also revealed an “obesity paradox” in which overweight individuals have a better short- and intermediate-term prognosis for survival when diagnosed with congestive heart failure (Horwich et al., 2018; Zamora et al., 2013), which may indicate some protective effect of fatness conferring survival in cattle experiencing cardiac remodeling.

Three separate estimates of ADG were calculated by creating an early window of ADG, a late window of ADG, as well as the full feeding period estimate of ADG using the weights at processing and harvest. Early ADG was estimated from processing to the first re-implant event, representing a window of early feedlot growth corresponding to the first 80 to 120 days on feed. Early ADG differed among heart score categories, with the fastest growing animals exhibiting a heart score of 2 or greater at harvest (P < 0.05). This supports an association between increased growth rate and cardiac remodeling during the feeding period. A longer window of ADG was also estimated representing the period from feedyard arrival to harvest. This longer period of ADG was also different among animals with different heart scores, with the fastest growing animals exhibiting a heart score of 2, 3, or 4 at harvest. The late ADG estimate represents the last 80 to 120 days on feed. For late ADG, individuals with a heart score of 5 had a significantly lower estimate of ADG (P < 0.05). This may suggest that the extreme level of cardiac insufficiency these animals are experiencing near harvest is impairing growth during the period leading up to harvest. Also, individuals with a heart score of 4 have similar growth rates as individuals with a heart score of 1, suggesting little to no impact on growth rate until cardiac remodeling reaches some threshold of severity.

Measurements for individual average DMI were collected at an average of 152 days on feed, and significantly differed among heart score categories. Individuals with a heart score of 5 had significantly higher DMI compared to individuals with a heart score of 1 (P < 0.05). A similar trend of increasing residual feed intake across heart scores was observed, with individuals with a heart score of 5 having a significantly higher residual feed intake compared to individuals with a heart score of 1 through 4. The association observed between cardiac remodeling and dry matter intake is supported by Heffernan et al. (2020) where a weak relationship between DMI, feed to gain, and heart score was noted. In this study, the larger sample size provides greater power for estimation of these effects.

Means and standard deviations for carcass traits observed for animals harvested under emergency harvest procedures with a heart score of 4 or 5 are displayed in Table 2. Emergency harvest shipments also include poor gaining animals, chronic lameness, and other chronic morbidity not necessarily associated with BCHF. In addition, 6.2% of the animals that were delivered to the packing plant using the emergency harvest procedures were condemned, indicating a higher level of generalized chronic morbidity. Approximately 29.61% of the 1,263 individuals harvested under emergency harvest criteria had a heart score 4 or 5, which indicates early shipment of suspected cardiac morbidities is an effective method to reduce feedyard death loss of animals susceptible to cardiac mortality and capture carcass value. It is important to note that the individuals in emergency shipments with a heart score of 1 through 3 were likely pulled early due to injury or morbidity not associated with cardiac remodeling. However, the lightest animals were still associated with a heart score 5, supporting the trend observed from the data for regular shipments. This relatively small population harvested in a highly variable time series generates the potential for sampling errors that make it difficult to draw conclusions from emergency harvest shipment phenotype data. Caution should be used when interpreting the observed trends, and additional investigation into population trends is warranted. In cases where a feedlot is in close proximity to a harvest facility, emergency harvest procedures may present a reasonable opportunity to reduce the economic losses associated with cardiac mortality while simultaneously improving animal welfare by reducing mortalities in the feedlot.

**Table 2.** Mean and standard deviation for carcass traits observed from animals harvested under emergency harvest procedures with heart score 4 or 5.

| Trait<sup>1</sup> | Heart Score |  |  |  |
| --- | --- | --- | --- | --- |
|  | 4 |  | 5 |  |
| | $\mu$ | $\sigma$ | $\mu$ | $\sigma$ |
| Harvest Wt (kg) | 551.39 | 79.84 | 531.50 | 74.23 |
| HCW (kg) | 345.46 | 53.95 | 325.33 | 45.34 |
| REA (cm<sup>2</sup>) | 32.87 | 4.52 | 30.58 | 4.21 |
| FAT (cm) | 1.34 | 0.62 | 1.34 | 0.59 |
| MARB | 474.56 | 101.16 | 497.59 | 118.33 |
| YG | 2.97 | 0.72 | 3.14 | 0.77 |
| N | 145 |  | 229 |  |
| % of Total Fragile | 11.48% |  | 18.13% |  |
| N Condemned | 7 |  | 35 |  |

<sup>1</sup>HCW = hot carcass weight, REA = ribeye area, FAT = backfat thickness, MARB = marbling score, YG = yield grade

Average genomic breed percentage for observed breed types with approximately known genetic background are displayed in Table 3. Individuals were categorized into the observed breed types based on visual appearance at feedlot processing in combination with known origin (ranch or dairy origin). On average, the genomic breed percentage estimate appears to be within approximately 5% of what the expected percentage would be based on origin, known breed types of sire and dam, and physical appearance. Given a robust reference panel covering major breed types, Low-Pass sequencing followed by imputation appears to provide an accurate platform for determining breed type and percentage from genotype calls (Snelling et al., 2020). Figure 2 displays a PCA plot using individual animal genotype for the 5,001 individuals in the reference population. This displays the variety of crossbred individuals with diverse genetic backgrounds available in the reference population, with the major breed types represented including Angus, Hereford, Charolais, Holstein, and Jersey. Figure 3 displays the same PCA analysis as Figure 2, but with BCHF cases highlighted in red. Cases of BCHF were observed in every breed class, but this figure displays a higher concentration of BCHF cases in breed classes mostly composed of Angus and Angus crossbreds.

**Table 3.** Average genomic breed percentage for major breed types from genotyped individuals with known genetic background.

| Observed Breed Type | Average Genomic Breed Percentage |  |  |  |  |
| --- | --- | --- | --- | --- | --- |
|  | Angus | Hereford | Charolais | Holstein | Other |
| Angus | 81.32% | 8.90% | 1.73% | 1.98% | 6.07% |
| Angus x Hereford | 46.81% | 46.30% | 0.96% | 1.56% | 4.37% |
| Angus x Dairy | 49.67% | 0.00% | 0.00% | 47.03% | 3.30% |
| Charolais x Dairy | 0.00% | 0.00% | 50.15% | 46.62% | 2.77% |
| Charolais x Angus x Hereford | 31.83% | 11.50% | 48.38% | 0.00% | 8.29% |

**Figure 2.**
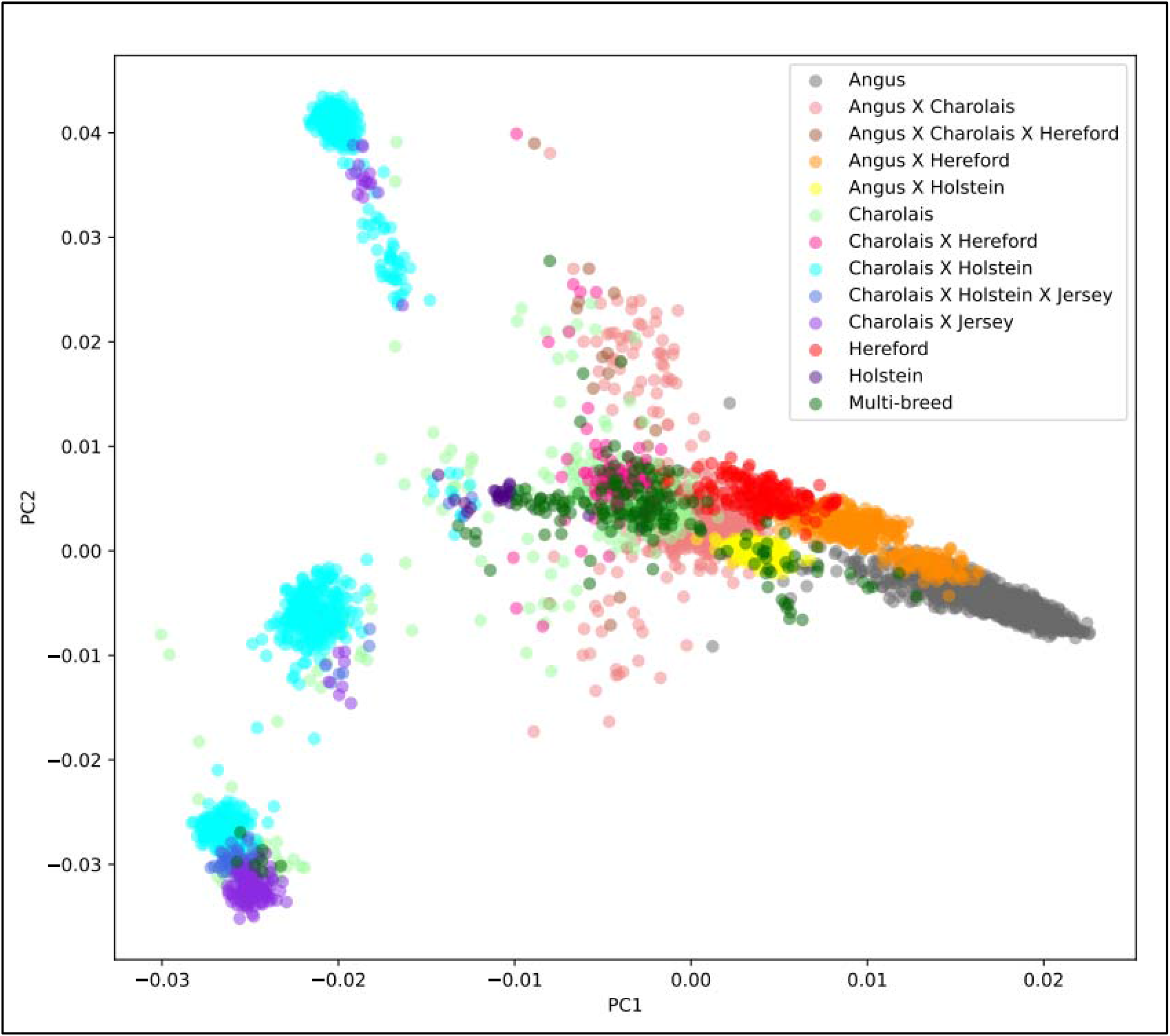
Principal component analysis (PCA) of 5,001 individuals with a Low-Pass genotype and a heart score phenotype colored by breed type determined by genomic breed percentage.igure 2

**Figure 3.**
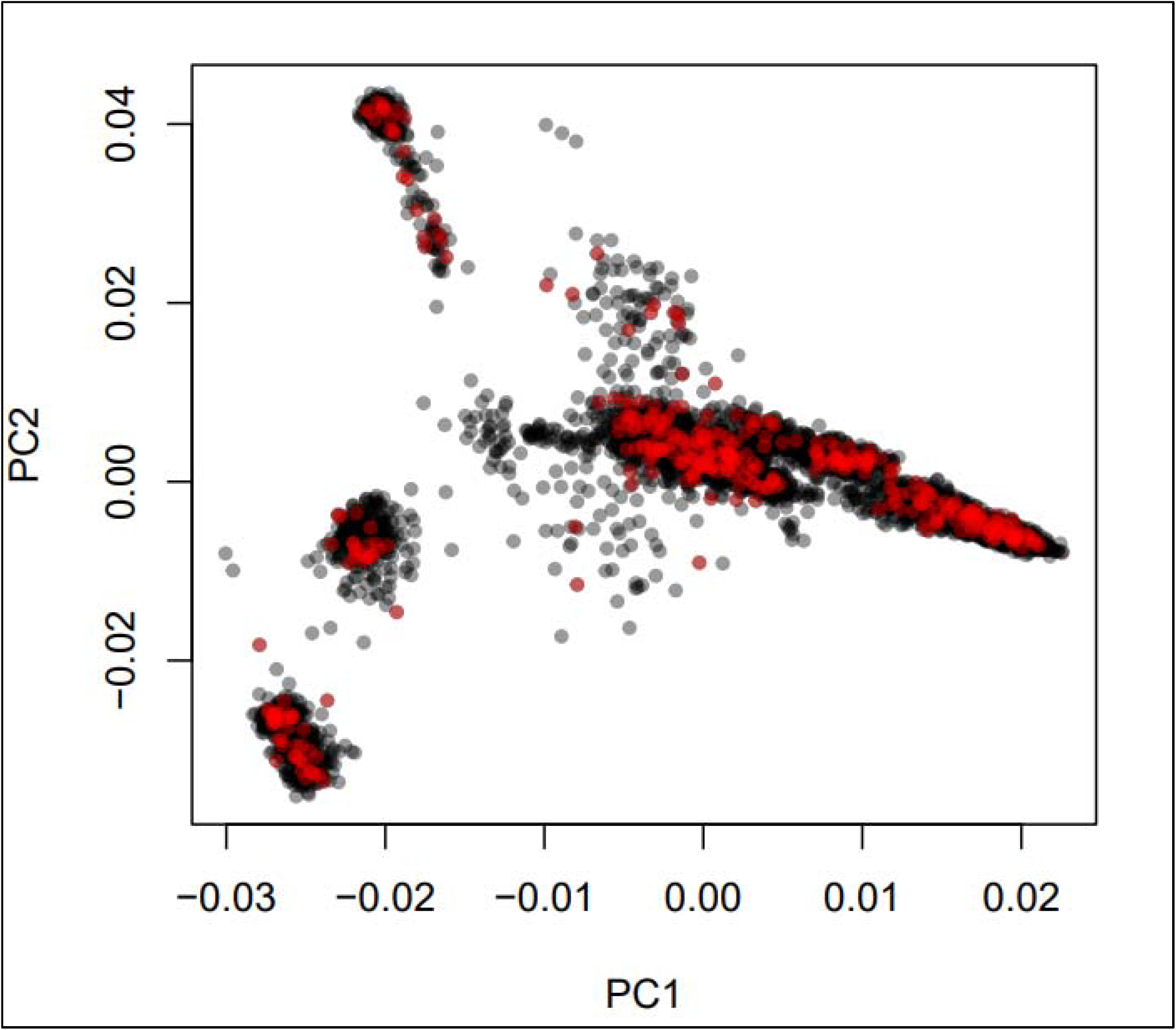
Principal component analysis (PCA) of 5,001 individuals with a Low-Pass genotype and a heart score phenotype colored by BCHF case (red) or normal (black) heart phenotype.

Distribution of heart score by observed breed type is displayed in Table 4. Individuals characterized as Angus breed type displayed the highest percentage of heart score 4 or 5 of any breed type observed. Individuals classified as Charolais x Angus x Hereford crosses had the lowest observed rate of heart score 4 or 5 at 1.42%. This observation potentially indicates that heterosis or dilution of the Angus breed direct effects through crossbreeding may result in a more normal cardiac morphology observed at harvest. Crosses using traditional beef breeds on dams of dairy origin also result in a lower risk of a heart score 4 or 5 with Angus x dairy and Charolais x dairy at 1.44% and 1.64%, respectively.

**Table 4.** Percent distribution and head count (in parentheses) of heart score by breed type according to observed phenotype and known background according to cattle origin.

| Observed Breed Type | Heart Score |  |  |  |  |
| --- | --- | --- | --- | --- | --- |
|  | 1 | 2 | 3 | 4 | 5 |
| Angus | 56.71%<br>(1,335) | 27.70%<br>(652) | 11.09%<br>(261) | 3.31%<br>(78) | 1.19%<br>(28) |
| Angus x Hereford | 62.85%<br>(1,308) | 24.94%<br>(519) | 9.18%<br>(191) | 1.92%<br>(40) | 1.11%<br>(23) |
| Angus x Dairy | 71.83%<br>(2,012) | 20.56%<br>(576) | 5.96%<br>(167) | 1.21%<br>(34) | 0.43%<br>(12) |
| Charolais x Dairy | 63.99%<br>(3,007) | 26.54%<br>(1,247) | 7.83%<br>(368) | 1.43%<br>(67) | 0.21%<br>(10) |
| Charolais x Angus x Hereford | 68.05%<br>(2,151) | 21.92%<br>(693) | 8.54%<br>(270) | 1.20%<br>(38) | 0.22%<br>(9) |

Binary logistic regression of individual breed percentage estimate on heart score (1 or 2 vs. 4 or 5) is displayed in Figure 4. Breed differences support the hypothesis of a genetic component influencing heart score and clinical BCHF. As the percentage of Angus breed type increases, the relative risk of observing a heart score of 4 or 5 increases in an approximately linear function. At 100% Angus breed percentage the relative risk of observing a heart score of 4 or 5 approaches 15% in this model. In contrast, as percent Hereford, Charolais, or Holstein breed percentage increases, the relative risk of observing a heart score of 4 or 5 decreases. At 100% Charolais or Holstein breed percentage the estimated risk of observing a heart score of 4 or 5 is below 5%. At 100% Hereford breed percentage the relative risk of observing a heart score of 4 or 5 is approximately 7%. Moxley et al. (2019) and Grandin (2022) identified BCHF as a problem of Angus cattle. The observed heart scores and breed characterization analysis in this population supports these literature findings.

**Figure 4.**
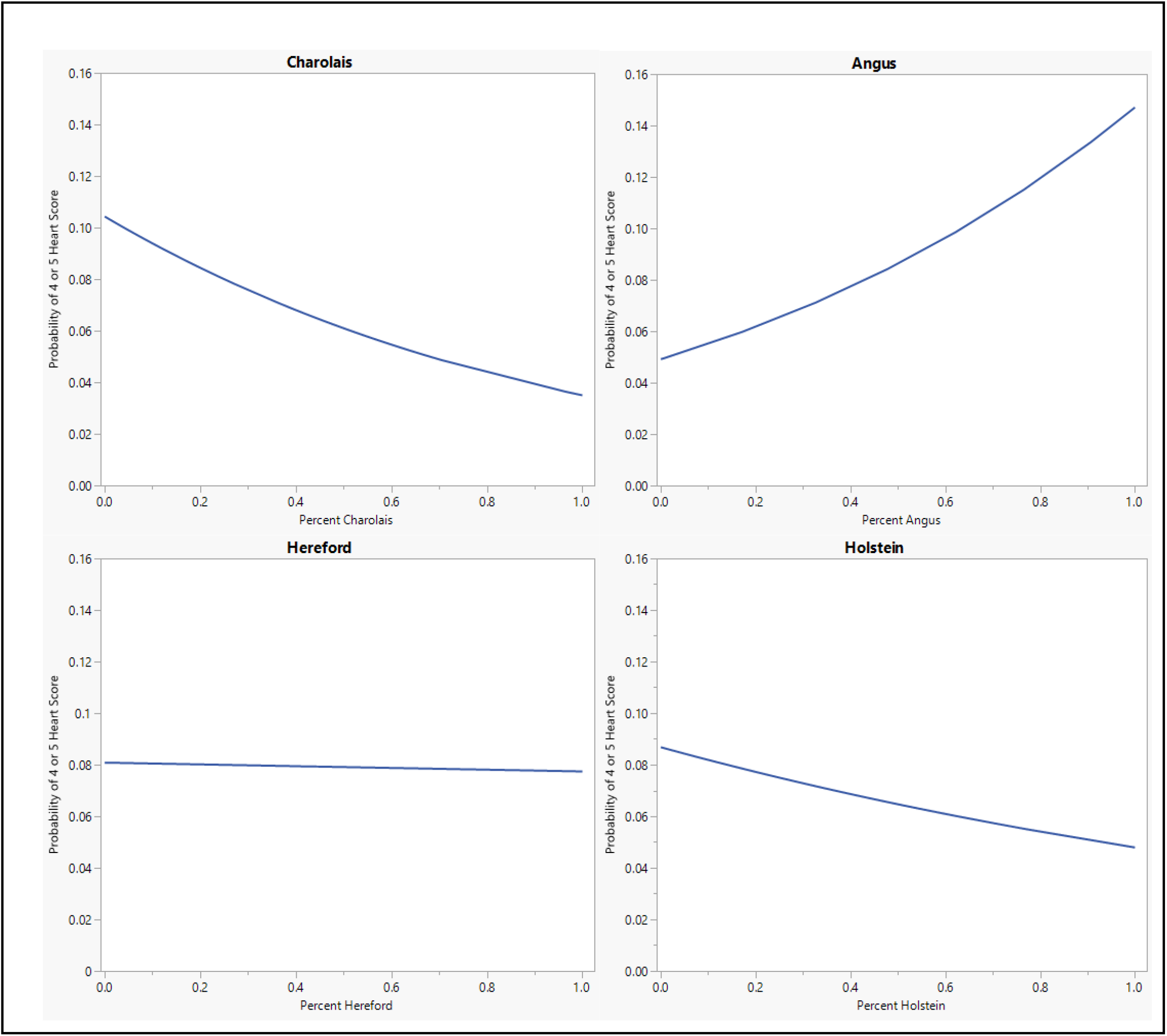
Binary logistic regression of individual genomic breed percentage estimate on heart score (1 or 2 vs. 4 or 5) for 4 major breed categories observed in this population.

Heaton et al. (2022) previously identified two genetic variants that were associated with BCHF. However, the lack of significant quantitative trait loci detected in a genome-wide association analysis in the present study (result not shown) indicates heart score and BCHF are less likely to be the result of a major gene effect and more likely to be polygenic traits. Mean and 95^th^ percentile estimates of variance components for heart score are displayed in Table 5. The heritability estimate of heart score modeled as a binary trait in this population was 0.356. To the extent that breed direct effects exist for BCHF and are heritable, this estimate contains those effects in addition to the usual within-breed additive genetic variation. Thus, heart score appears to be moderately heritable when modeled as a binary (BCHF case vs. control) phenotype in a multi-breed population. Genetic correlations between heart score and additional production traits estimated from a two-trait binary-threshold model are displayed in Table 5. A moderate and positive genetic correlation was observed between heart score and traits related to feed intake and growth. The highest genetic correlation was observed with HCW at 0.460, which indicates that individuals with a genetic potential to reach higher live weight and carcass weight at harvest are more likely to develop a heart score of 4 or 5 at harvest. Similarly, a genetic correlation of 0.289 was observed between binary heart score and ADG, which supports the association between growth rate and risk of abnormal cardiac morphology.

**Table 5.** Mean and 95^th^ percentile of variance components estimates for bovine congestive heart failure (BCHF) as well as genetic correlations from a threshold linear model of BCHF and production traits.

| Component<sup>1</sup> | Binary Threshold-Linear Gibbs Model |  |  |
| --- | --- | --- | --- |
|  | 2.5 Percentile | Mean | 97.5 Percentile |
| $\sigma_g^2$ BCHF | 0.105 | 0.554 | 0.914 |
| $\sigma_e^2$ BCHF | 0.942 | 1.003 | 1.062 |
| $h^2$ BCHF | 0.100 | 0.356 | 0.463 |
| $r_g$ BCHF - ADG | -0.038 | 0.289 | 0.441 |
| $r_g$ BCHF - DMI | -0.065 | 0.309 | 0.477 |
| $r_g$ BCHF - HCW | 0.358 | 0.460 | 0.515 |
| $r_g$ BCHF – REA | -0.688 | 0.080 | 0.277 |
| $r_g$ BCHF – MARB | -0.622 | -0.108 | 0.139 |
| $r_g$ BCHF - FAT | -0.359 | -0.120 | 0.054 |

<sup>1</sup> $\sigma_g^2$ = genetic variance, $\sigma_e^2$ = residual variance, $h^2$ = heritability, $r_g$ = genetic correlation

The binary expression of BCHF could be used to create an EPD for use in selection programs to reduce the incidence of individuals developing severe cardiac remodeling prior to harvest over time. To determine the ability to predict trait differences from genomic data, the accuracy of EPD prediction was estimated using a GBLUP model. First using 4,639 individuals (373 cases and 4,266 controls), the GBLUP model was fit and used to predict 362 individuals with known phenotypes that were harvested in another season (182 cases and 180 controls) and found Pearson’s correlation of 0.241 between EPD and true case/control status. Both data sets were then compiled to produce a new data set with 5,001 individuals (555 cases, and 4,446 controls). This new data set was used to perform a 5-fold cross-validation with the GBLUP model. In this test, an average Pearson correlation of 0.344 was observed in the test splits. These results indicate that an EPD for BCHF is a feasible solution for reducing the prevalence of this condition through genomic selection, and additional data collection efforts would improve the accuracy of this EPD over time.

## 4 Conclusions

Congestive heart failure is a growing concern for multiple segments of the beef industry and a selection tool is needed to consider this trait in breeding objectives for beef cattle. In addition, reducing the incidence of cardiac remodeling and heart failure would result in an industry-wide improvement in animal welfare and reduced mortality. It is important to note that the mortality rate reported in this study is likely lower than the average rate across the industry due to the proximity of this particular feedlot to a packing plant. Emergency harvest procedures alone are an ineffective mitigation strategy for the average feedlot due to the distance required to transport cattle to a harvest facility. Heart score observed at harvest is a moderately heritable trait and has significant genetic correlation to carcass and growth traits. A significant association between BCHF and Angus or Angus crossbred cattle is also apparent in this study, which may require many operations across multiple sectors of the industry to weigh the risk associated with elevated death loss against any production related characteristics associated with breed selection. This study also indicates an EPD for heart score could be implemented as a selection tool to reduce the incidence of this trait over time. There is a need for evaluating how this trait fits into modern selection indexes given the significant genetic correlation to other economic traits under selection in cattle. If ignored, the incidence of congestive heart failure is suspected increase over time given the direction of the genetic correlation to other economically important traits that carry a high value in modern indexes and are subsequently under strong selection.

## 6 Author Contributions

RCR, JWB, JLH, LEF, and MDM conceptualized the study and contributed to data analysis and manuscript editing. JWB and LEF authored the manuscript. JKP, LEF, and JLH created and analyzed low-pass genotypes. ARN, RCR, and JWB collected and analyzed the phenotype data.

## 8 Acknowledgments

The authors would like to acknowledge Tammy Oltman, Naya Bradshaw, Vicky Herrera Vargas, Joel Ontiveros, and Tommie Crockett for the collection of heart scores and phenotype data reported in this study.

## 9 Availability of Data Statement

The data analyzed in this study is owned by Simplot Livestock Co. Requests to access the data can be directed to: Randall Raymond.

